# Social contact behaviors are associated with infection status for whipworm (*Trichuris* sp.) in wild vervet monkeys (*Chlorocebus pygerythrus*)

**DOI:** 10.1101/2020.10.07.329409

**Authors:** Brandi Wren, Ian S. Ray, Melissa Remis, Thomas R. Gillespie, Joseph Camp

## Abstract

Social grooming in the animal kingdom is common and serves several functions, from removing ectoparasites to maintaining social bonds between conspecifics. We examined whether time spent grooming with others in a highly social mammal species was associated with infection status for gastrointestinal parasites. Of six parasites detected, one (*Trichuris* sp.) was associated with social grooming behaviors, but more specifically with direct physical contact with others. Individuals infected with *Trichuris* sp. spent significantly less time grooming conspecifics than those not infected, and time in direct contact with others was the major predictor of infection status. One model correctly predicted infection status for *Trichuris* sp. with a reliability of 95.17% overall when the variables used were time spent in direct contact and time spent grooming others. This decrease in time spent grooming and interacting with others is likely a sickness behavior displayed by individuals with less energy or motivation for non-essential behaviors. This study highlights the need for an understanding of a study population’s parasitic infections when attempting to interpret animal behavior.

## Introduction

Grooming is widespread in the animal kingdom, from insects [1-4] to rodents [5-8], birds [9-13], and primates [14-17]. Grooming is generally classified into two types: self-grooming, in which individuals groom themselves, and social grooming or allogrooming in which individuals groom others or are groomed by others [18]. Both self-grooming and social grooming have hygienic benefits including removal of parasites and debris [19-21], as well as physiological benefits like releasing endorphins and lowering heart rate [22,23]. Social grooming specifically also has social, reproductive, and resource acquisition benefits [24-28]. Grooming with others can also be costly, however, because it may lead to transmission of viruses, bacteria, and other pathogens [29-34]. Studies of rodents [35] and primates [36,37] have even suggested that social grooming may increase likelihood of infection with parasites that are not typically transmitted between hosts.

Grooming and sociality in general have an influence on reproductive fitness. For example, social integration (often measured largely by grooming behaviors) was associated with increased reproductive success in feral mares [38]. Social integration was also positively correlated with survival of infants of female savannah baboons (*Papio cynocephalus*) [39]. Allopreening was related to an increase in reproductive fitness in common guillemots (*Uria aalge*) [40].

Grooming and social contact are known to provide health benefits as well. When horses are groomed on favored body parts, they exhibit reduced heart rates [23]. In a case study of a pigtail macaque (*Macaca* sp.), being groomed by others resulted in a significantly lower heart rate but self-grooming and initiating grooming with others did not [41]. In rhesus macaques (*Macaca mulatta*), being groomed mitigated anxiety regarding a dominant conspecific as revealed by faster heart rate deceleration when being groomed [42]. It is common for individuals to become so relaxed that they fall asleep when being groomed [personal observations]. A study on Barbary macaques (*Macaca sylvanus*) suggested that grooming others can reduce stress, as indicated by assessments of fecal glucocorticoids [43].

Research supports claims that social grooming can help reduce numbers of ectoparasites, often termed the hygiene hypothesis [14]. In one study of Japanese macaques, researchers concluded that the main function of grooming was to eliminate ectoparasites, specifically lice [19]. Another study revealed that wild savannah baboons (*Papio cynocephalus*) that were groomed more frequently had fewer ticks [44]. Further, research suggests that social grooming may reduce mortality risk from the fungus *Metarhizium anisopliae* among termite hosts (*Zootermopsis angusticollis*) [45].

However, this approach does not fully acknowledge the complexity of all host-parasite relationships and the multifaceted relationship between grooming and health. Understanding host-parasite ecology means understanding the complex interplay of a number of factors including distribution of parasites in the environment and likelihood of encountering them, age, sex, physiology, and social behavior [46,47]. Dunbar [25] concluded that the hygiene hypothesis alone could not account for primate social grooming behaviors because a meta-analysis revealed that amount of time spent grooming with others correlated more with group size than with body size across the Order Primates overall. Dunbar’s work suggested that social grooming may serve more of a hygienic function among New World monkeys (among whom grooming time correlated more precisely with body weight than group size) while serving more of a social function among Old World monkeys and apes (among whom grooming time correlated more precisely with group size than body weight). Further, some studies have found that parasite loads do not correlate with grooming behaviors. For example, grooming behaviors in the Seychelles warbler (*Acrocephalus sechellensis*) were not correlated with feather mite load [48]. Observers also noted that chacma baboons (*Papio ursinus*) did not always remove ticks (*Rhipicephalus* sp.) from partners when grooming with them, even when researchers could see from afar that ticks were engorged [49]. It is also worth noting that tick infestations were estimated to cause over half of known infant deaths among that study population.

Some parasites may be more likely to be transmitted when two individuals groom, and it is not uncommon for some types of pathogens, like viruses, to be transmitted through social contact. This is being increasingly acknowledged as societies around the world have dealt with the COVID-19 pandemic, and researchers have noted these connections between social proximity and the spread of infectious disease in both humans and other animal species [50]. Social contact in western lowland gorillas (*Gorilla gorilla gorilla*) – largely observed as grooming – was associated with death from the Ebola-Zaire virus in a study population in Congo [30]. One study on meerkats (*Suricata suricatta*) revealed that those who groomed others more frequently were more likely to become infected with tuberculosis (*Mycobacterium bovis*) than those that groomed others less frequently [51]. Social grooming in ants (*Lasius* sp.) resulted in transmission of the potentially pathogenic fungus *Metarhizium anisopliae*, however this ultimately aids in developing immunity to the fungus [52]. Examples like this highlight the complexity of the host-parasite relationship and the need for a more nuanced approach to understanding host-parasite dynamics.

Some studies even suggest that grooming may play a role in transmission of macroparasites like nematodes which are not typically considered to be capable of host-to-host transmission. For example, grooming among mice (*Mus* sp.) was associated with infection with the gastrointestinal parasite *Heligmosomoides polygyrus* [35]. Grooming behaviors in vervet monkeys (*Chlorocebus aethiops [pygerythrus]*) also varied with infection status with hookworm species [36].

Behavioral ecology theory predicts that an adaptation is selected for when the costs of a behavior are exceeded by the benefits. Social grooming presents significant benefits including removal of ectoparasites and debris, increased likelihood of future social support or tolerance, increased likelihood of future mating opportunities, and improved reproductive fitness. However, social grooming also presents significant costs (e.g., increased likelihood of contracting an infectious pathogen leading to death) because of the close contact. While much is known about the benefits of social grooming, much less is known about the costs of it among nonhuman primates.

The aim of this study was to determine whether grooming behaviors in a highly social mammal species varied with respect to infection status with gastrointestinal parasites. We examined various dimensions of vervet monkey (*Chlorocebus pygerythrus*) grooming behavior, including time spent grooming others, time spent being groomed by others, and time spent in direct contact with others. We tested fecal samples for gastrointestinal parasites, specifically protozoa and helminths. We then tested whether individuals who were infected with parasites spent similar relative amounts of time grooming and/or receiving grooming from other individuals. We predicted that if social grooming or direct social contact facilitates the transmission of any gastrointestinal parasite species in our study population, then those individuals that spend more time grooming with others should be more likely to exhibit infection with parasites.

## Materials and methods

### Study site and subjects

Data were collected from three social groups of wild vervet monkeys (*Chlorocebus pygerythrus*) at Loskop Dam Nature Reserve (LDNR), South Africa (Figure 1). LDNR is located in the Olifants River Valley within Mpumalanga and Limpopo provinces (25°25’S, 29°18’E), and is managed by the Mpumalanga Tourism and Parks Agency (MTPA). The reserve is 225 km^2^ and surrounds Lake Loskop, a reservoir of 23.5 km^2^. The reserve encompasses both highveld and bushveld ecological zones, and habitat ranges from open grasslands to dense woodlands [53]. Common woody species throughout the three groups’ ranges include a variety of species of *Combretum, Acacia, Rhus, Grewia*, and *Ficus*, as well as *Dichrostachys cinerea, Mimusops zeyheri*, and *Olea europa* [54,55]. Altitude in LDNR ranges from 990-1450 m and the reserve exhibits a highly seasonal climate. Annual rainfall during the study was 914.5 mm, most of which fell between October and January [2009-2010, LDNR, unpublished. *data*]. Average minimum and maximum temperatures during the study period were 13.5°C and 26.1°C, respectively [2009-2010, LDNR unpublished. *data*; 2009-2010, Wren unpublished. *data*].

**Figure 1:**
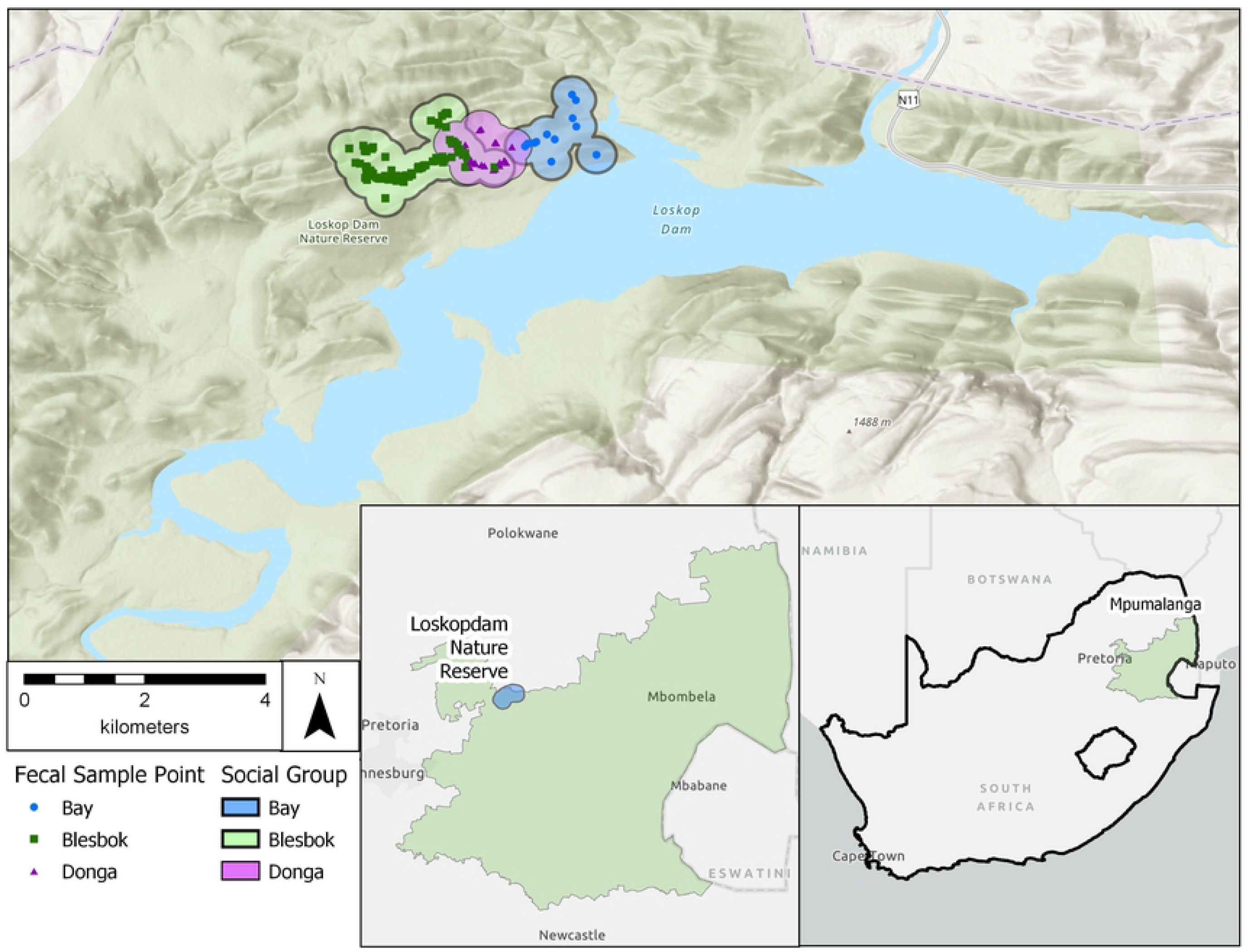
Loskop Dam Nature Reserve, South Africa.

We chose *Chlorocebus pygerythrus* as the study species because individuals exhibit variation in grooming behaviors [56], allowing us to examine differences in the relationship between social behaviors and parasite infection status. Groups of *Ch. pygerythrus* in LDNR – and much of the surrounding region – typically vary in size from 13-25 individuals [54,55,57]. Six groups of *Ch. pygerythrus* at LDNR are habituated, and researchers have been conducting studies of these groups semi-regularly for more than a decade [58-61]. We collected data from three of the six habituated groups at LDNR: Blesbok group, Donga group, and Bay group. At the commencement of the study there were 14 individuals in the Blesbok group, 16 in the Donga group, and 17 in the Bay group; the total study population fluctuated due to births, migrations, and deaths, and was 54 at the conclusion of the study. Here we present data on a total of 55 subjects as well as a subset of 38 of those study subjects. Information on group composition for each social group can be found in Wren [55] and Wren et al. [36,62]. We located groups using known sleeping sites and home ranges. Data were recorded for only the Blesbok group from July – October 2009 because other researchers were studying the Donga and Bay groups during that time. Data were collected from all three social groups for the remainder of the project.

### Behavioral data collection

From July 2009 through July 2010 we followed *Ch. pygerythrus* groups and collected observational data [63]. Although most data were collected between 07:00 h and 16:00 h, observations were conducted between 05:00 h and 19:00 h. We performed 30-min focal follows on each individual. Some follows were terminated early because monkeys were lost from sight, but we kept data from all follows longer than 5 min. We used continuous recording for all behavioral data [63].

We recorded data on the following variables for all bouts of social grooming: start and stop times, whether grooming was given or received, and identity of grooming partner. A grooming bout was considered to be a new bout when any of the following conditions were met: the focal individual stopped grooming or being groomed for 30 seconds or more, the direction of grooming switched (i.e., the individual being groomed began grooming its partner or vice versa), or the individual switched grooming partners. We also recorded data on start and stop times for direct physical contact with another individual and identification of direct social contact partners.

### Fecal sample collection and analysis

We collected 332 fecal samples non-invasively from identified individuals directly following defecation, and samples were immediately preserved in a 10% buffered formalin solution. We recorded data on the following variables for each sample: date, individual, social group, location (GPS coordinates), consistency and color of feces, and whether adult worms were visible in the stool. Fecal samples varied from approximately 3 to 7 g.

We used three methods to detect parasite eggs and cysts in samples in order to reduce the risk of false negatives: fecal flotation, fecal sedimentation, and immunofluorescence microscopy. We isolated helminth eggs and protozoan cysts and oocysts from fecal material using fecal flotation with double centrifugation (at 1800 rpm for 10 min) in NaNO_3_ solution and fecal sedimentation with dilute soapy water [64]. We also used immunofluorescence microscopy with a Merifluor *Cryptosporidium/Giardia* Direct Immunofluorescent Detection Kit (Meridian Bioscience Inc., Cincinnati, Ohio, USA) to detect *Cryptosporidium* sp. oocysts and *Giardia* sp. cysts [65]. Parasite eggs and cysts were identified by egg or cyst shape, size, color, and contents for flotation and sedimentations, and measurements of eggs and cysts were taken with an ocular micrometer fitted to a compound microscope. For the immunofluorescence microscopy, we scored fecal samples for the presence or absence of *Cryptosporidium* sp. oocysts and *Giardia* sp. cysts.

### Data analysis

Although we collected 511 h of behavioral data and 332 fecal samples from 55 individual vervet monkeys, we analyzed 477.66 h of data and 272 fecal samples from 38 individuals. Some behavioral data were not used in the final analyses because there were no corresponding fecal samples for some study subjects, and vice versa.

We calculated measures of parasite infections following Bush et al. [66]. Richness refers to the number of parasite species detected in a host or group. Prevalence refers to the number of hosts infected with a specific parasite species divided by the number of hosts examined.

We used analysis of variance (ANOVA) to examine whether social groups differed by sampling effort, grooming behaviors, or parasite infections. We planned contrasts to compare each group and each combined grouping of groups to identify statistically significant differences in each observed variable. We further explored significant results from the ANOVA using a logistic regression model from calculated z-scores for each independent variable. We set the presence of *Trichuris* sp. as the binary outcome variable with the following observed variables: time observed, total seconds observed, number of grooming partners, number of grooming partners giving grooming, number of grooming partners receiving grooming, number of total contact partners, time spent giving grooming, time spent receiving grooming, time spent self grooming, time in direct contact, and time spent playing.

We used GNU PSPP 1.2.0 for all statistical tests. We set the significance level at *p < 0*.*05* and considered all tests two-tailed.

### Ethical note

This study was conducted with approvals from LDNR, MTPA, Applied Behavioural Ecology and Ecosystem Research Unit of the University of South Africa, and Purdue University’s Animal Care & Use Committee (approval #07-609). We followed all guidelines for the study of nonhuman primates set forth by the International Primatological Society.

## Results

### Descriptive statistics

Mean time observed per individual for the entire study sample was 9.3 hours (*n* = 55, minimum = 0.18, maximum = 37.04, SD = 8.84). Age of individuals ranged from 1 to 11 years (*n* = 55, mean = 4.80, SD = 2.67) at the end of the study, and 42% of subjects were female (35/55) while 64% were male (35/55).

For parasitological hypothesis testing, we used a subset of the entire study sample that consisted of 38 individuals from the three social groups. This included only individuals for which both behavioral and parasitological data were available and complete. Mean time observed per individual in this subset used for parasitological hypothesis testing was 12.57 hours (*n* = 38, minimum = 1.343, maximum = 37.04, SD = 8.83).

Mean number of fecal samples collected per individual was 7.16 (*n* = 38, minimum = 1, maximum = 25, SD = 8.84). For this subset of individuals for which parasitological results were obtained, groups differed significantly in regard to total time observed (F_(2,35)_ = 28.242, *p* < 0.001) and number of fecal samples collected (F_(2,35)_ = 27.929, *p* < 0.001). Age of individuals ranged from 3 to 11 years (*n* = 38, mean = 5.6, SD = 2.29) at the end of the study, and 42% of subjects were female (16/38) while 58% were male (22/38).

### Behavioral results

Mean proportion of time spent grooming others was 5.0% of total time observed (*n* = 55, mean = 0.05, minimum = 0.0, maximum = 0.29, SD = 0.06). (Table 1) Mean proportion of time spent being groomed for the entire study sample was 4.0% of total time observed (*n* = 55, mean = 0.04, minimum = 0.0, maximum = 0.01, SD = 0.03). (Table 2) Social groups did not differ in regard to mean time spent grooming others (F_(2,52)_ = 0.39, *p* = 0.677) or being groomed by others (F_(2,52)_ = 0.93 *p* = 0.401). Time in direct contact, while not grooming, accounted for an average of 20% of the total time observed and did not differ among groups (F_(2,52)_ = 0.03, *p* = 0.971). For the subset of the study sample used in the parasitological hypothesis testing, mean proportion of time spent being groomed was also 4.0% of total time observed (*n* = 38, mean = 0.04, minimum = 0.0, maximum = 0.01, SD = 0.025). Mean proportion of time spent grooming others was 5.1% of total time observed (*n* = 38, mean = 0.05, minimum = 0.0, maximum = 0.29, SD = 0.059). Social groups did not differ in regard to mean time spent grooming others (F_(2,35)_ = 0.172, *p* > 0.05) or being groomed by others (F_(2,35)_ = 0.517, *p* > 0.05). Groups also did not differ in mean time spent in direct contact (F_(2,35)_ = 2.6, *p* > 0.05).

**Table 1:** Mean percentage of time spent grooming others among vervet monkeys (*Chlorocebus pygerythrus*) at Loskop Dam Nature Reserve, South Africa

|  | Infected, Mean<br>Time Observed | Not Infected,<br>Mean Time<br>Observed | t-value | P-value |
| --- | --- | --- | --- | --- |
| *Trichuris* | 4% | 14% | -3.17 | 0.003 |
| Hookworm | 6% | 4% | 0.87 | 0.388 |
| Spirurids | 5% | 6% | -0.90 | 0.375 |
| *Oesophagostomum* sp. | 4% | 9% | -1.74 | 0.090 |
| *Strongyloides* sp. | 3% | 6% | -1.45 | 0.156 |
| *Entamoeba coli* | 3% | 5% | -0.76 | 0.454 |

Degrees of freedom = 36 in all cases.

**Table 2:** Mean percentage of time spent being groomed among vervet monkeys (*Chlorocebus pygerythrus*) at Loskop Dam Nature Reserve, South Africa

|  | Infected,<br>Mean Time<br>Observed | Not Infected,<br>Mean Time<br>Observed | t-value | P-value |
| --- | --- | --- | --- | --- |
| *Trichuris* sp. | 4% | 4% | 0.02 | 0.987 |
| Hookworm | 4% | 4% | 0.80 | 0.430 |
| Spirurids | 4% | 4% | -0.39 | 0.698 |
| *Oesophagostomum*<br>sp. | 4% | 5% | -0.69 | 0.496 |
| *Strongyloides* sp. | 4% | 4% | -0.44 | 0.660 |
| *Entamoeba coli* | 3% | 4% | -0.50 | 0.622 |

Degrees of freedom = 36 in all cases.

### Parasitological results

Analyses revealed six types of parasites: *Trichuris* sp. (92% prevalence in the study sample), hookworm (71% prevalence), spirurids (68% prevalence), *Oesophagostomum* sp. (84% prevalence), *Strongyloides* sp. (24% prevalence), and *Entamoeba coli* (92% prevalence) (*n* = 38). Descriptions of these can be found in Wren et al. [35,53,60]. We did not detect presence of *Cryptosporidium* sp. or *Giardia* sp. with immunofluorescence microscopy.

Social group differences are presented in Table 3. These differences are likely due to the different sampling efforts for each social group as noted in the methods section. Social groups were significantly different regarding richness of parasite species detected (F_(2,35)_ = 5.0804, *p* = 0.012). The Bay group differed significantly from the Donga group (*t*(35) = 2.98, *p* = 0.005) and the Blesbok from the Donga group (*t*(35) = 2.58, *p* = 0.014). The combination of the Bay and Blesbok group differed significantly from the Donga group (*t*(35) = 2.05, *p* = 0.048) while the combination of the Blesbok and Donga group differed significantly from the Bay group (*t*(35) = 3.16, *p* = 0.003).

**Table 3:** Levene’s Test of Homogeneity of Variance for Behavioural Variables in Vervet Monkeys (*Chlorocebus pygerythrus*) at Loskop Dam Nature Reserve, South Africa

| Variable | Levene's<br>Statistic | df1 | df2 | *p* |
| --- | --- | --- | --- | --- |
| Total Seconds Observed | 6.27 | 2 | 52 | 0.004 |
| Number of Grooming Partners | 5.52 | 2 | 52 | 0.007 |
| Number of Grooming Partners (Giving) | 4.89 | 2 | 52 | 0.011 |
| Number of Grooming Partners (Receiving) | 3.16 | 2 | 52 | 0.051 |
| Number of Total Partners (Contact) | 3.81 | 2 | 52 | 0.029 |
| Time Giving Grooming | 2.68 | 2 | 52 | 0.078 |
| Time Receiving Grooming | 2.90 | 2 | 52 | 0.064 |
| Time Self-Grooming | 8.80 | 2 | 52 | 0.001 |
| Time in Direct Contact | 0.28 | 2 | 52 | 0.756 |
| Time Playing | 4.79 | 2 | 52 | 0.012 |

Groups differed significantly in the presence of hookworm (F_(2,35)_ = 1.35, *p* = 0.272), The Bay group differed from the Blesbok group (*t*(35) = 3.45, *p* = 0.001) and the Donga group (*t*(35) = 2.44, *p* = 0.020). The combination of the Bay and Donga groups differed from the Blesbok group (*t*(35) = 2.65, *p* = 0.012). The combination of the Blesbok and Donga group differed significantly from the Bay group (*t*(35) = 3.34, *p* = 0.002).

Groups did not differ significantly in the presence of *Trichuris* sp. (F_(2,35)_ = 1.35, *p* = 0.272), spirurids (F_(2,35)_ = 2.80, *p* = 0.075), *Oesophagostomum* sp. (F_(2,35)_ = 0.91, *p* = 0.412), *Strongyloides* sp. (F_(2,35)_ = 0.17, *p* = 0.842), or *Entamoeba coli* (F_(2,35)_ = 1.35, *p* = 0.272).

### Hypothesis testing results

#### Group differences

One-way analysis of variance (ANOVA) was conducted to examine group differences in observed behaviors. Levene’s test of homogeneity of variance revealed only the number of grooming partners, time receiving grooming, and time in direct contact met this assumption (Table 3). However, because ANOVA is robust with respect to violations of homogeneity of variance analyses could still be performed. There were statistically significant differences among groups for: total seconds observed (F_(2, 52)_ = 22.79, *p* < 0.001); number of grooming partners (F_(2, 52)_ = 15.70, *p* < 0.001); number of grooming partners giving (F_(2, 52)_ = 8.11, *p* = 0.001); number of total partners (F_(2, 52)_ = 19.08, *p* < 0.001); time self-grooming (F_(2, 52)_ = 3.54, *p* = 0.036) (Table 4).

**Table 4:**
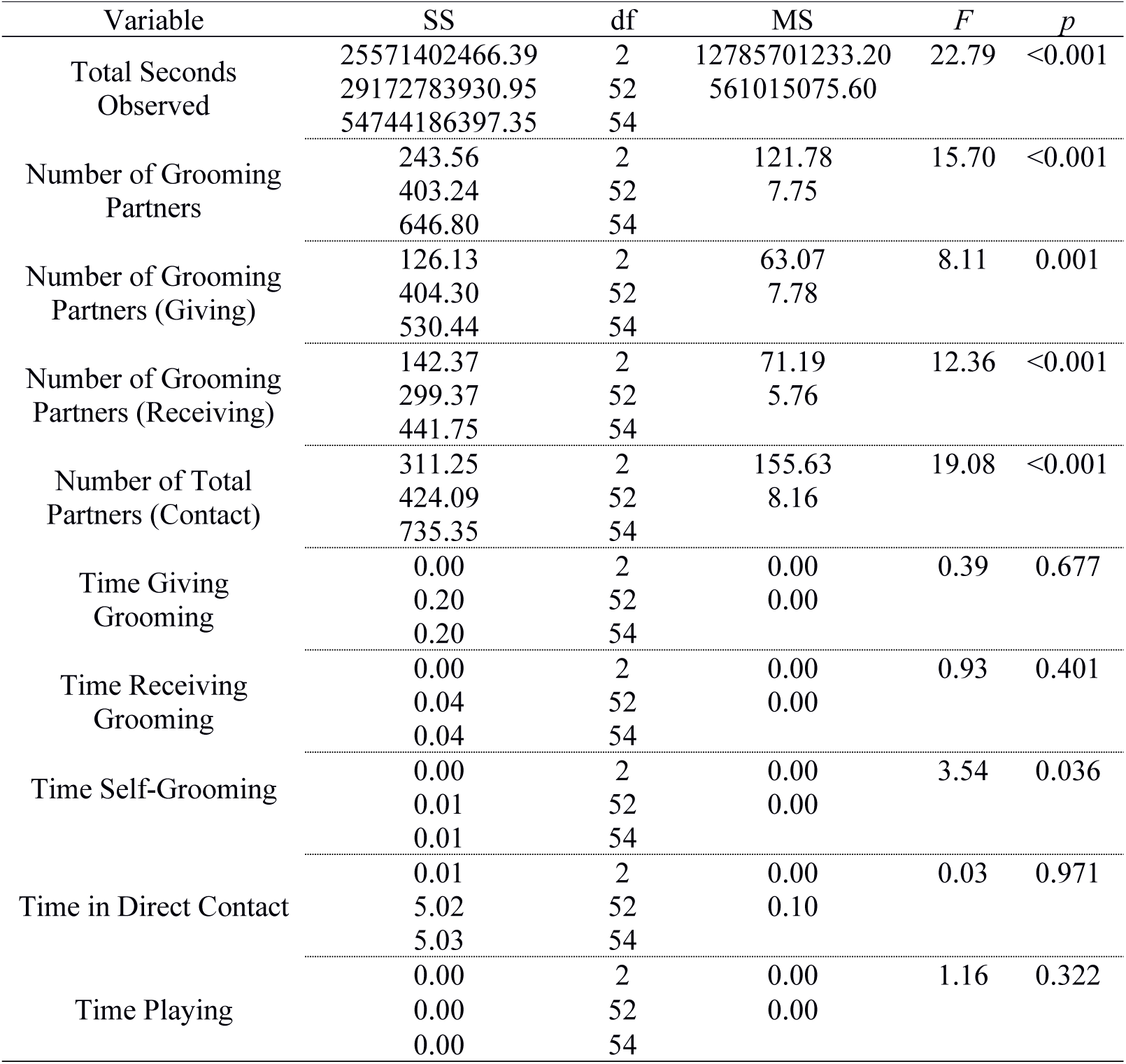
ANOVA Summary for Behavioural Variables in Vervet Monkeys (*Chlorocebus pygerythrus*) at Loskop Dam Nature Reserve, South Africa

Planned contrasts revealed specific differences among groups and combinations of groups (Tables 5 & 6). There were statistically significant differences for all combinations of groups with respect to total seconds observed. There were statistically significant differences for all combination of groups with respect to the number of grooming partners except between the combination of the Bay and Blesbok groups compared to the Donga group, (*t*(22.12) = 0.86, *p* = 0.398). There were statistically significant differences for all combination of groups with respect to the number of grooming partners giving grooming except between the combination of the Bay and Blesbok groups compared to the Donga group, (*t*(36.79) = 0.63, *p* = 0.543). There were statistically significant differences for all combination of groups with respect to the number of grooming partners receiving grooming except between the combination of the Bay and Blesbok groups compared to the Donga group, (*t*(52) = 0.1.03, *p* = 0.310). There were statistically significant differences for all combination of groups with respect to the number of total partners except between the combination of the Bay and Blesbok groups compared to the Donga group, (*t*(34.10) = 1.21, *p* = 0.236). There were no statistically significant differences for any combination of groups with respect to time giving grooming. There were no statistically significant differences for any combination of groups with respect to time receiving grooming. There were statistically significant differences for all combinations of groups with respect to time spent self-grooming except for: the Bay group compared to the Donga group, (*t*(32.88) = 0.54, *p* = 0.59); the combination of the Bay and Blesbok groups compared to the Donga group, (*t*(32.62) = −0.79, *p* = 0.435); the Bay group compared to the combination of the Blesbok and Donga groups, (*t*(23.03) = −1.60, *p* = 0.123). There were no statistically significant differences for any combination of groups with respect to the time spent playing except between the combination of the Bay and Blesbok groups compared to the Donga group, (*t*(28.23) = 2.07, *p* = 0.048).

**Table 5:** Contrast Coefficients for ANOVA of Behavioural Variables in Vervet Monkeys (*Chlorocebus pygerythrus*) at Loskop Dam Nature Reserve, South Africa

| Contrast | Group |  |  |
| --- | --- | --- | --- |
|  | 1<br>(Bay) | 2<br>(Blesbok) | 3<br>(Donga) |
| 1 | 1 | 0 | -1 |
| 2 | 1 | -1 | 0 |
| 3 | 0 | 1 | -1 |
| 4 | 1 | 1 | -2 |
| 5 | 1 | -2 | 1 |
| 6 | -2 | 1 | 1 |

**Table 6:**
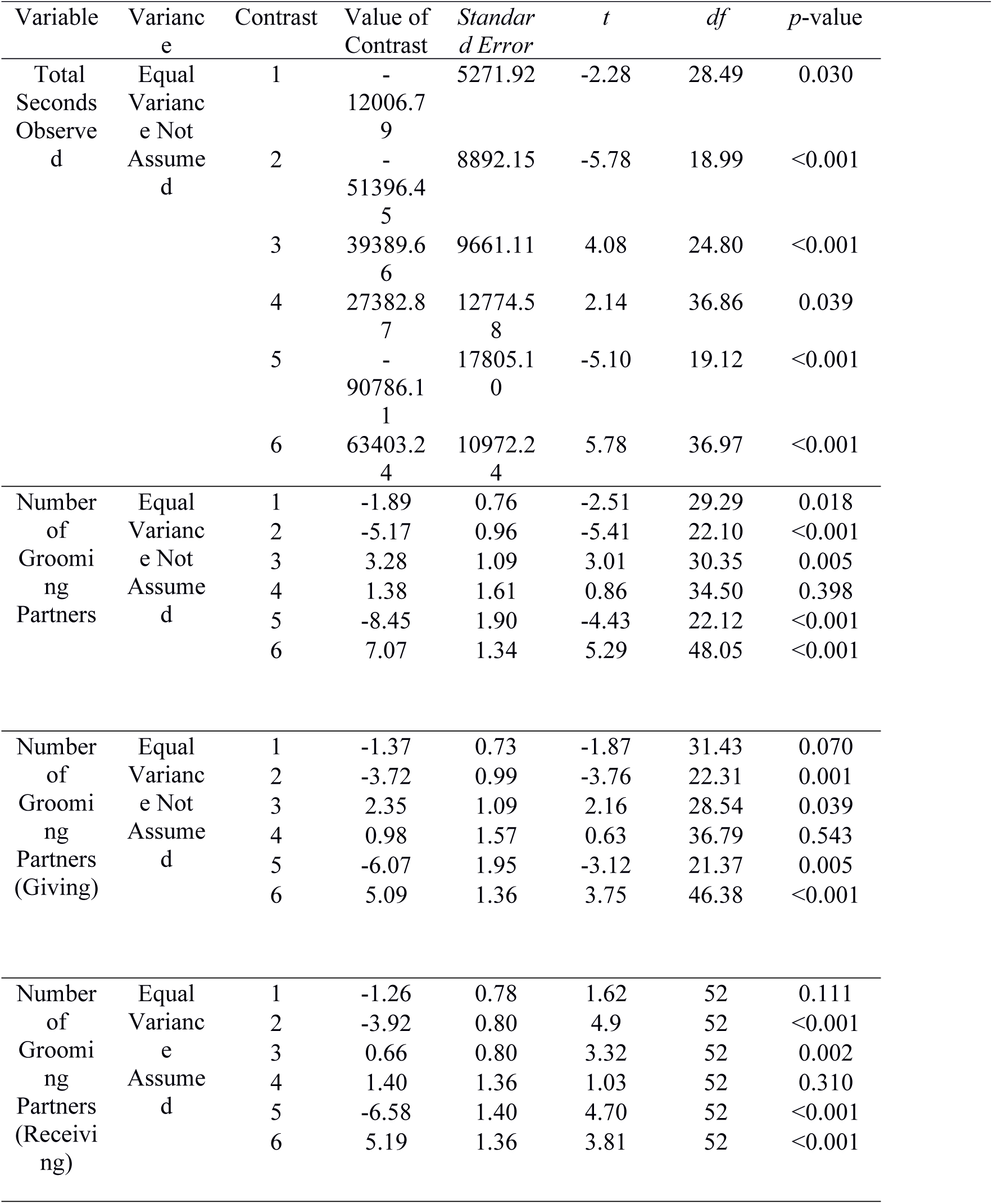
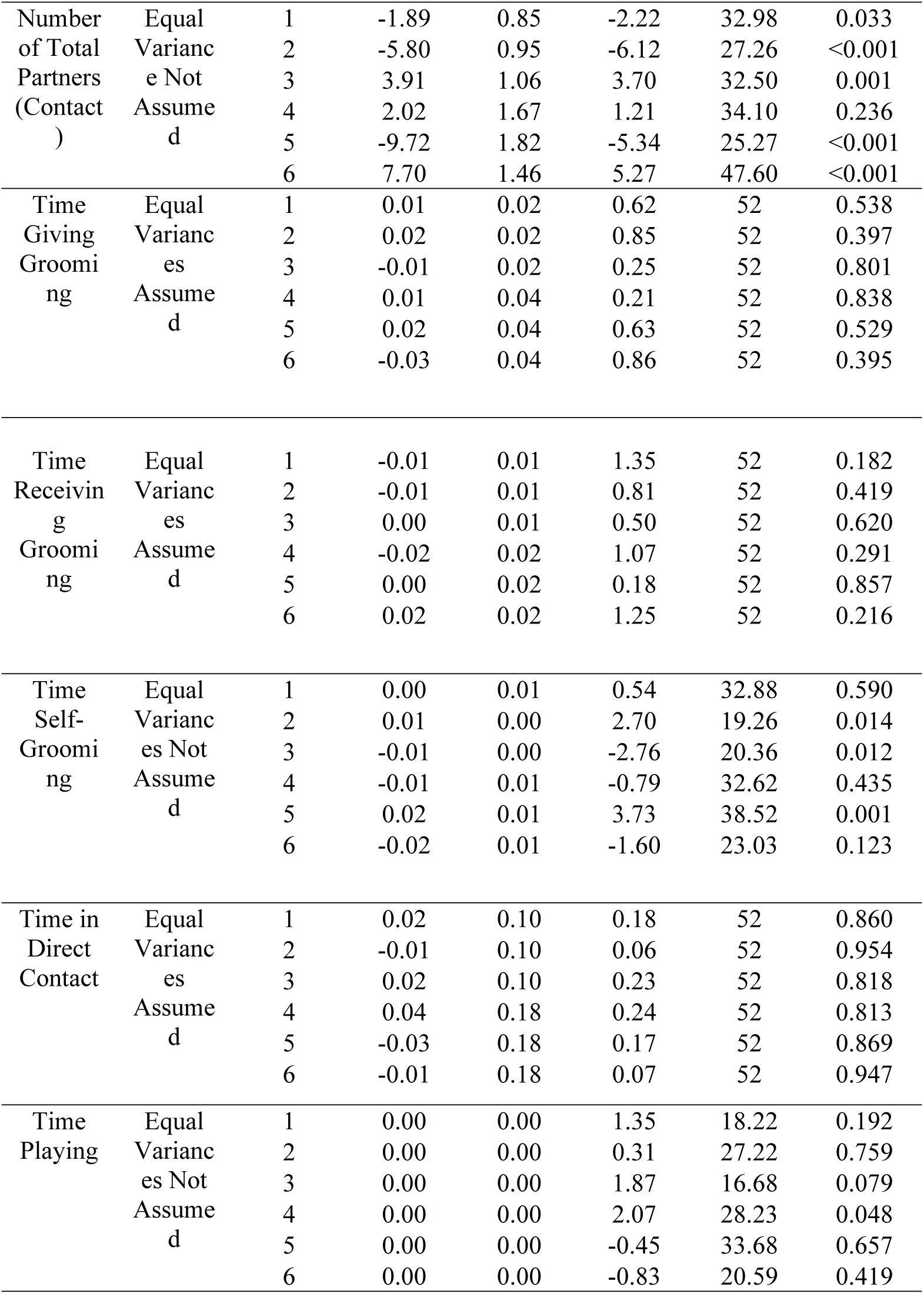
ANOVA Follow-up Results for Behavioural Variables in Vervet Monkeys *hlorocebus pygerythrus*) at Loskop Dam Nature Reserve, South Africa

### Generalized linear model

All behavioral variables from all observed individuals were converted to z-scores for inclusion in the generation of a multivariate logistic regression equation, with the dependent variable modeled as the presence or absence of *Trichuris* sp. The only individual statistically significant effect was time in direct contact (β = −4.63, *p* = 0.02; Table 7). The model shows excellent fit based on the −2 Log likelihood (24.22) and Nagelkerke pseudo-R^2^ (0.73) (Table 8). This equation was able to correctly predict 100% of observed absence of *Trichuris* sp. and 94.44% of observed presence of *Trichuris* sp., for a total of 95.17% of cases correctly predicted (Table 9).

**Table 7:**
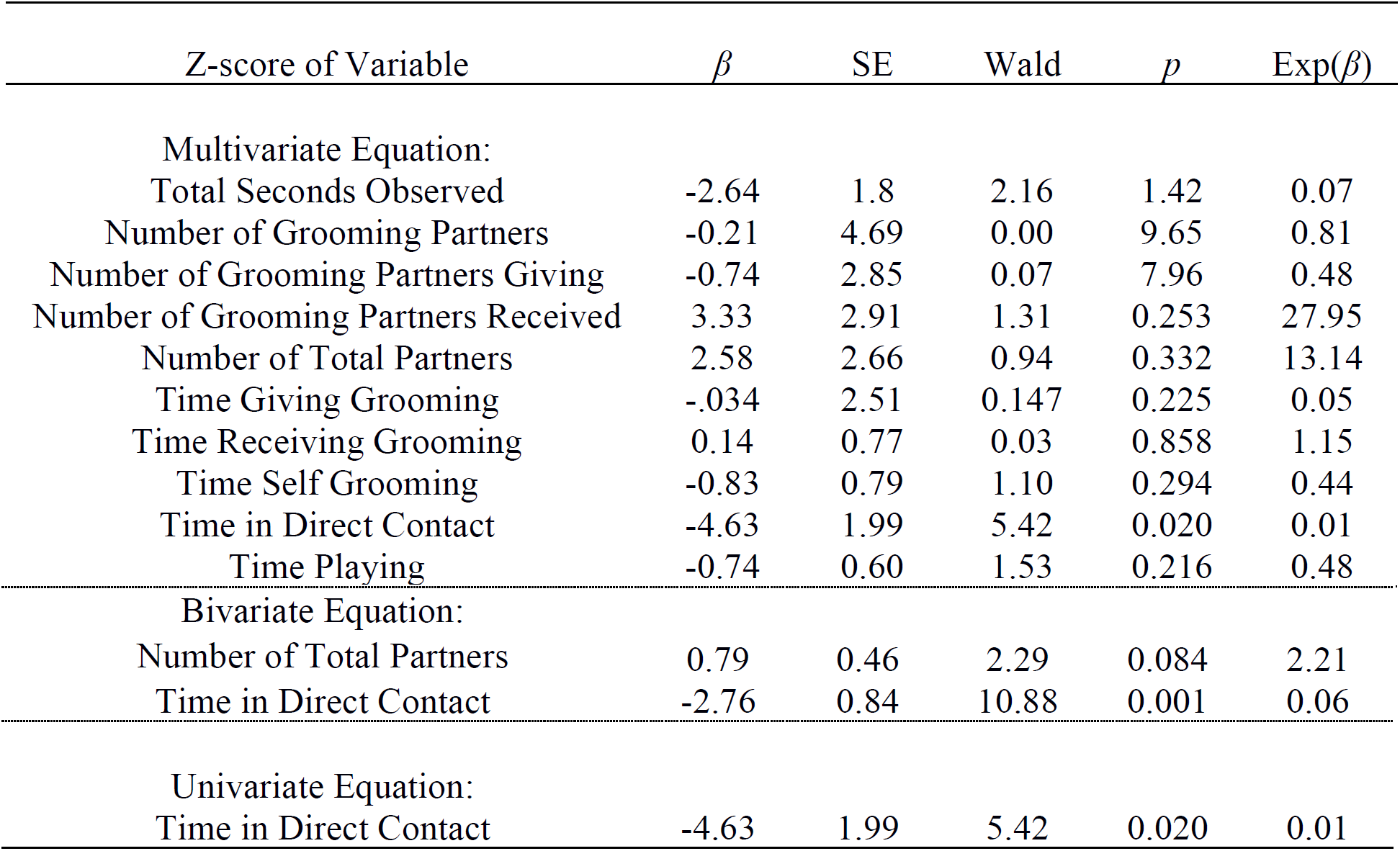
Logistic Regression Summary Table for Univariate and Equations Predicting *Trichuris* infection status in Vervet Monkeys (*Chlorocebus pygerythrus*) at Loskop Dam Nature Reserve, South Africa

**Table 8:**
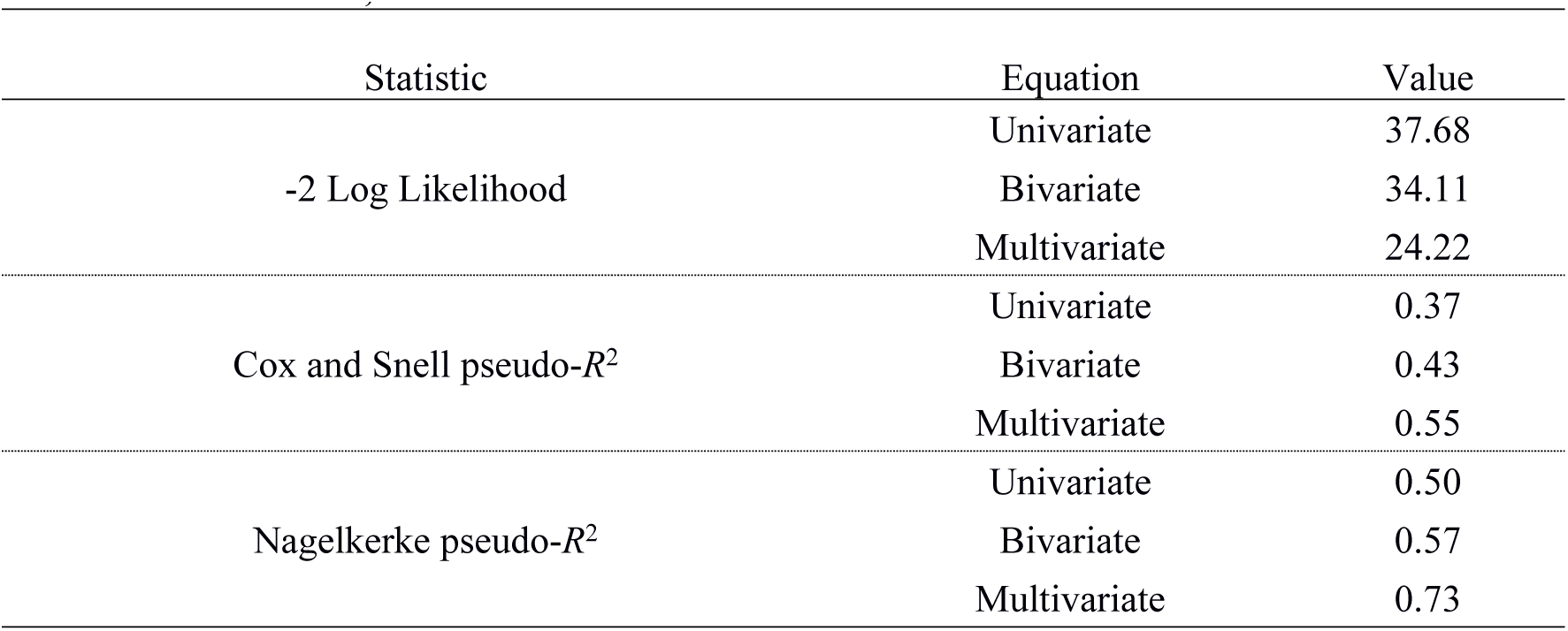
Logistic Regression Fit Statistics for Univariate and Multivariate Equations edicting *Trichuris* infection status in Vervet Monkeys (*Chlorocebus pygerythrus*) at Loskop m Nature Reserve, South Africa

**Table 9:**
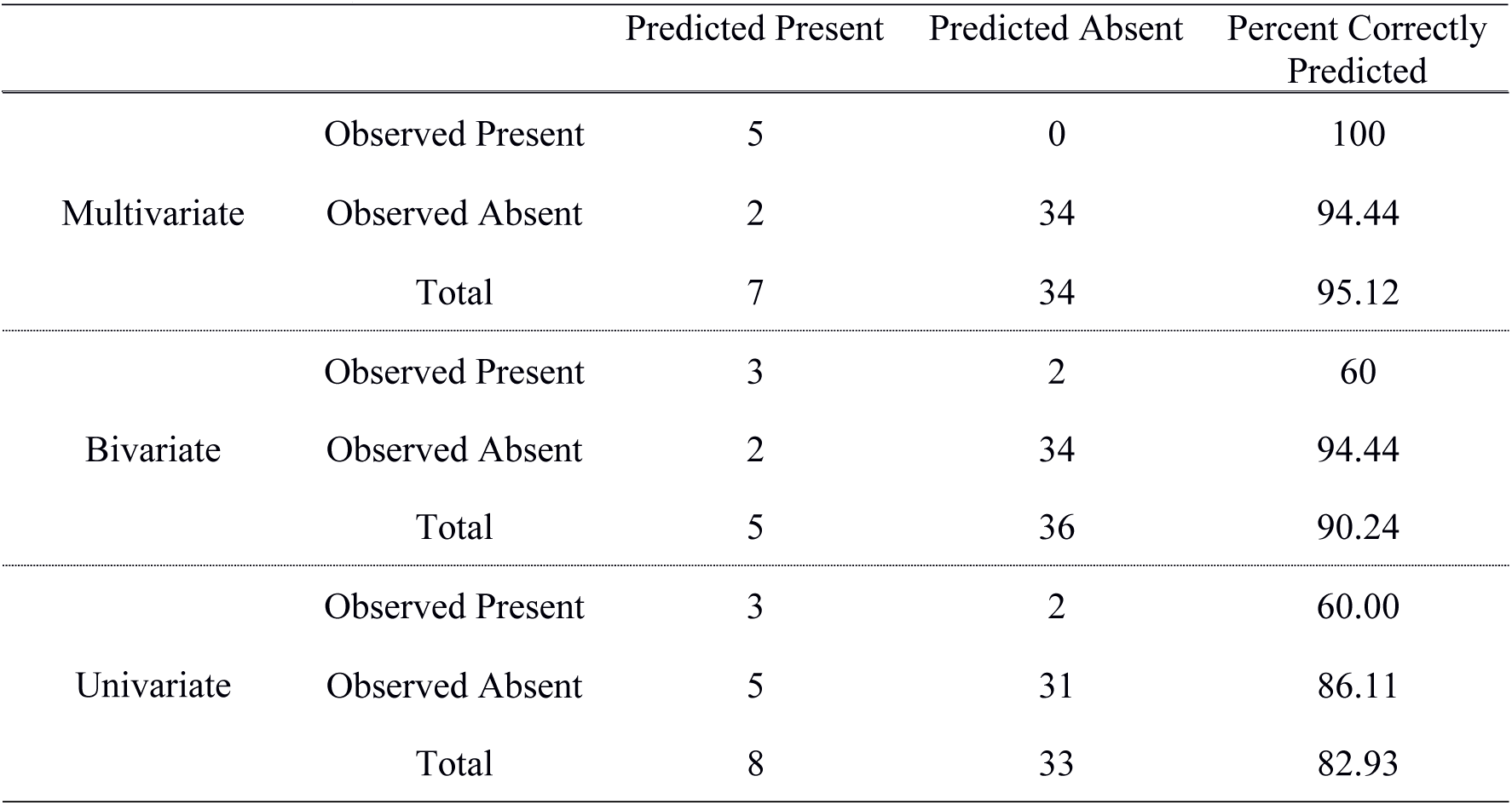
Logistic Regression Classification by Univariate and Multivariate Equations for predicting *Trichuris* infection status in Vervet Monkeys (*Chlorocebus pygerythrus*) at Loskop m Nature Reserve, South Africa

|  | Predicted Present | Predicted Absent | Percent Correctly Predicted |
| --- | --- | --- | --- |

|  |  |  |  |  |
| --- | --- | --- | --- | --- |
|  | Observed Present | 5 | 0 | 100 |
| Multivariate | Observed Absent | 2 | 34 | 94.44 |
|  | Total | 7 | 34 | 95.12 |
|  | Observed Present | 3 | 2 | 60 |
| Bivariate | Observed Absent | 2 | 34 | 94.44 |
|  | Total | 5 | 36 | 90.24 |
|  | Observed Present | 3 | 2 | 60.00 |
| Univariate | Observed Absent | 5 | 31 | 86.11 |
|  | Total | 8 | 33 | 82.93 |

A bivariate logistic regression was conducted using only the z-score of time in direct contact and the total number of grooming partners to predict the presence of *Trichuris* sp. This regression indicated that time in direct contact with others had a statistically significant negative effect on infection status (β = −2.76, *p* = 0.001). Total number of grooming partners had a non-statistically significant positive effect on infection status (β = 0.79, *p* = 0.084). The model fit for this equation is worse than for the multivariate equation, based on a higher −2 Log likelihood (34.11) and lower Nagelkerke pseudo-R^2^ (0.73) (Table 8). This equation was able to correctly predict 60% of observed absence of *Trichuris* sp. and 94.44% of observed presence of *Trichuris* sp., for a total of 90.24% of cases correctly predicted (Table 9).

A univariate logistic regression was conducted using only the z-score of time in direct contact and the presence of *Trichuris* sp. This regression indicated that time in direct contact with others had a statistically significant effect on infection status (β = −2.73, *p* = 0.001). The model fit for this equation is worse than for the multivariate equation, based on a higher −2 Log likelihood (37.68) and lower Nagelkerke pseudo-R^2^ (0.50) (Table 8). This equation was able to correctly predict 60% of observed absence of *Trichuris* sp. and 86.11% of observed presence of *Trichuris* sp., for a total of 82.93% of cases correctly predicted (Table 9).

## Discussion

Vervet monkeys at LDNR that are infected with *Trichuris* sp. tend to spend significantly less time grooming conspecifics as well as less time in direct contact with others when compared to those that are not infected. Infected individuals at LDNR spent an average of 4% of their observed time grooming others, while those not infected with this parasite spent an average of 14% of their observed time grooming others. However, no differences existed in time spent being groomed by others. Overall, for the entire sample (*n* = 55), study subjects spent about 20% of their time in direct contact with another individual. The subset used for parasitological analysis spent 6.5% (*n* = 38) of their time in direct contact with another individual. This large difference is primarily influenced by the inclusion of infants and mothers with infants in the entire sample of *n* = 55, but only mothers in the smaller subset of *n* = 38. These mother-infant dyads remain in almost constant contact for the first weeks of a monkey’s life and this inflates the overall mean for the group. Because there were not enough fecal samples from these infants, their behavioral data was not included in hypothesis testing. One model that we built was able to correctly predict infection status for *Trichuris* sp. with a reliability of 95.17% overall, but the major factor for prediction was time spent in direct contact.

These results do not support the hypothesis that social grooming facilitates transmission of this type of gastrointestinal parasite. One possible explanation for these results is that individuals that are infected with *Trichuris* sp. experience degraded health and/or less motivation to groom others and interact with others. Red colobus monkeys (*Procolobus rufomitratus*) in Uganda that were infected with *Trichuris* sp. decreased their time spent performing a number of behaviors, including grooming others [67]. Those same individuals spent more time resting as well as ingesting plant species and/or parts that suggest self-medicative behavior. Whipworm is known to cause anemia, chronic dysentery, rectal prolapse, and poor growth in humans with symptomatic infections [68], so less energy, motivation, or interest for behaviors like social grooming should not be surprising in other species.

Another possible explanation is that *Trichuris* sp. more directly alters host behavior in vervet monkeys. Gastrointestinal parasites are known to alter host behavior in some host-parasite relationships, an idea referred to as the manipulation hypothesis [69-72]. For example, *Toxoplasma gondii* causes intermediate rodent hosts to be more attracted to the scent of felid predators, which are also the definitive host for the parasite [73]. *Dicrocoelium dendriticum* causes infected ants to wait on the tips of blades of grass where they can be ingested by sheep, the parasite’s definitive host. Because manipulation of host behavior usually serves to facilitate transmission of the parasite from an intermediate host to a definitive host, and vervet monkeys do not serve as intermediate hosts for *Trichuris* sp., the manipulation hypothesis does not adequately explain the results of this study.

Other studies have found multiple morphotypes of *Trichuris* sp. in nonhuman primate hosts in captivity in Nigeria [74,75], suggesting that potentially multiple species of *Trichuris* sp. may infect nonhuman primates. The major implication of this has been seen as relevant for public health because it may mean that the species of *Trichuris* sp. that infect humans and nonhuman primates are not the same, suggesting that transmission of *Trichuris* sp. between humans and other primates is not as severe a public health concern as previously considered. However, it could also have implications for how primate hosts respond to or become infected with *Trichuris* sp.

Hart [76] noted that ill or infected animals display altered behavior, and argued that these sickness behaviors can be adaptive. One study of chimpanzees (*Pan troglodytes schweinfurthii*) revealed that infected individuals exhibit altered behavior, most fittingly described as lethargy [77]. Behavioral changes due to parasitic infections in fish have been observed and range from mating behaviors to foraging efficiency (reviewed in Barber et al. [78]). The Ghai et al. [67] study that revealed that *Trichuris* sp. was associated with a reduction in grooming and mating and also found that individuals infected with this parasite took longer to switch behaviors than those individuals that were not infected. These studies make it increasingly clear that host-parasite dynamics have far-reaching consequences for animal behavior.

This study suggests that the gastrointestinal parasite *Trichuris* sp. is associated with behavioral differences, specifically decreased time spent grooming others and time spent in direct contact with others, in vervet monkey hosts. These behavioral differences are extreme enough to influence group means when assessing behavior. Further, if an individual is less likely to groom or interact with conspecifics, then they may also experience lower social status and thus lower reproductive fitness. These results highlight the need for parasitological analyses for a complete and nuanced understanding of animal behavior.

## Acknowledgments

We would like to thank the Mpumalanga Parks and Tourism Agency, University of South Africa, and Loskop Dam Nature Reserve, specifically Leslie Brown, Tersia De Beer, and Jannie Coetzee, for research permissions and site support. We are also grateful to Katie Dean, Claire Detrich, Ruby Malzoni, Liz Sperling, Andrew Kalmbach, Lisa Breland, and others for assistance in the field as well as to Elizabeth Canfield, Naomi Hauser, and Michele Moses for assistance in the laboratory.

## References

1. Hlavac TF. Grooming systems of insects: structure, mechanics. Ann Entomol Soc Am. 1975;68:823–826.

2. Thomson JD. Pollen transport and deposition by bumble bees in Erythronium: influences of floral nectar and bee grooming. J Ecol. 1986;74:329–341.

3. Sumana A, Starks PT. Grooming patterns in the primitively eusocial wasp Polistes dominulus. Ethology. 2004;110:825–833.

4. Li C. A video-tracking method to identify and understand circadian patterns in Drosophila grooming [dissertation]. Coral Gables (FL): University of Miami; 2016.

5. Greer JM, Capecchi MR. Hoxb8 is required for normal grooming behaviour in mice. Neuron. 2002;33:23–34.

6. Ferkin MH, Leonard ST. Self-grooming by rodents in social and sexual contexts. Acta Zool Sinica. 2004;51:772–779.

7. Hawlena H, Bashary D, Abramsky Z, Krasnov BR. Benefits, Costs and constraints of anti-parasitic grooming in adult and juvenile rodents. Ethology. 2007;113:394–402.

8. Kalueff AV, Stewart AM, Song C, Berridge KC, Graybiel AM, Fentress JC. Neurobiology of rodent self-grooming and its value for translational neuroscience. Nat Rev Neurosci. 2016;17:45–59.

9. Clayton DH, Cotgreave P. Relationship of bill morphology to grooming behaviour in birds. Anim Behav. 1994;47:195–201.

10. Cotgreave P, Clayton DH. Comparative analysis of time spent grooming by birds in relation to parasite load. Behaviour. 1994;131:171–187.

11. Christe P, Richner H, Oppliger A. Of great tits and fleas: sleep baby sleep … Anim Behav. 1996;52:1087–1092.

12. Lenouvel P, Gomez D, Théry M, Kreutzer M. Do grooming behaviours affect visual properties of feathers in male domestic canaries, Serinus canaria? Anim Behav. 2009;77:1253–1260.

13. Villa SM, Goodman GB, Ruff JS, Clayton DH. Does allopreening control avian ectoparasites? Biol Letters. 2016;12:20160362.

14. Hutchins M, Barash DP. Grooming in primates: implications for its utilitarian function. Primates. 1976;17:145–150.

15. Seyfarth RM. The distribution of grooming and related behaviours among adult female vervet monkeys. Anim Behav. 1980;28:798–813.

16. Cheney DL. Intragroup cohesion and intergroup hostility: the relation between grooming distributions and intergroup competition among female primates. Behav Ecol. 1992;3:334–345.

17. Parr LA, Matheson MD, Bernstein IS, de Waal FB. Grooming down the hierarchy: allogrooming in captive brown capuchin monkeys, Cebus apella. Anim Behav. 1997;54:361–367.

18. Benjamin D. The development of grooming and its expression in adult animals. Ann NY Acad Sci 1988:525;1–7.

19. Tanaka I, Takefushi H. Elimination of external parasites (lice) is the primary function of grooming in free-ranging Japanese macaques. Anthropol Sci. 1993;101:187–193.

20. Böröczky K, Wada-Katsumata A, Batchelor D, Zhukovskaya M, Schal C. Insects groom their antennae to enhance olfactory acuity. P Natl Acad Sci. 2013;110:3615–3620.

21. Zhukovskaya M, Yanagawa A, Forschler BT. Grooming behaviour as a mechanism of insect disease defense. Insects. 2013;4:609–630.

22. Keverne EB, Martensz ND, Tuite B. Beta-endorphin concentrations in cerebrospinal fluid of monkeys are influenced by grooming relationships. Psychoneuroendocrino. 1989;14:155–161.

23. Feh C, de Mazières J. Grooming at a preferred site reduces heart rate in horses. Anim Behav. 1993;46:1191–1194.

24. Seyfarth RM. A model of social grooming among adult female monkeys. J Theor Biol. 1977;65:671–698.

25. Dunbar RI. Functional significance of social grooming in primates. Folia Primatol. 1991;57:121–131.

26. Henzi SP, Barrett L. The value of grooming to female primates. Primates. 1999;40:47–59.

27. Silk JB. Social components of fitness in primate groups. Science. 2007;317:1347–1351.

28. Schino G, Aureli F. Grooming reciprocation among female primates: a meta-analysis. Biol Letters. 2008;4:9–11.

29. Fromont E, Courchamp F, Pontier D, Artois M. Infection strategies of retroviruses and social grouping of domestic cats. Can J Zool. 1997;75:1994–2002.

30. Caillaud D, Levréro F, Cristescu R, Gatti S, Dewas M, Douadi M, Ménard N. Gorilla susceptibility to Ebola virus: the cost of sociality. Curr Biol. 2006;16:R489–R491.

31. Craft ME, Hawthorne PL, Packer C, Dobson AP. Dynamics of a multihost pathogen in a carnivore community. J Anim Ecol. 2008;77:1257–1264.

32. Drewe JA, Eames KT, Madden JR, Pearce GP. Integrating contact network structure into tuberculosis epidemiology in meerkats in South Africa: implications for control. Preventive Vet Med. 2011;101:113–120.

33. Rushmore J, Caillaud D, Matamba L, Stumpf RM, Borgatti SP, Altizer S. Social network analysis of wild chimpanzees provides insights for predicting infectious disease risk. J Anim Ecol. 2013;82:976–986.

34. Rushmore J, Bisanzio D, Gillespie TR. Making connections: insights from primate-parasite networks. Trends Parasitol. 2017;33:547–560.

35. Hernandez AD, Sukhdeo MV. Host grooming and the transmission strategy of Heligmosomoides polygyrus. J Parasitol, 1995;865–869.

36. Wren BT, Remis MJ, Camp JW, Gillespie TR. Number of Grooming Partners Is Associated with Hookworm Infection in Wild Vervet Monkeys (Chlorocebus pygerythrus). Folia Primatol. 2016;87:168–179.

37. Rimbach R, Bisanzio D, Galvis N, Link A, Di Fiore A, Gillespie TR. Brown spider monkeys (Ateles hybridus): a model for differentiating the role of social networks and physical contact on parasite transmission dynamics. Philos T R Soc B. 2015;370(1669):20140110.

38. Cameron EZ, Setsaas TH, Linklater WL. Social bonds between unrelated females increase reproductive success in feral horses. P Natl Acad Sci. 2009;106:13850–13853.

39. Silk JB, Alberts SC, Altmann J. Social bonds of female baboons enhance infant survival. Science. 2003;302:1231–1234.

40. Lewis S, Roberts G, Harris MP, Prigmore C, Wanless S. Fitness increases with partner and neighbour allopreening. Biol Letters. 2007;3:386–389.

41. Boccia ML, Reite M, Laudenslager M. On the physiology of grooming in a pigtail macaque. Physiol Behav. 1989;45:667–670.

42. Aureli F, Preston SD, de Waal F. Heart rate responses to social interactions in free-moving rhesus macaques (Macaca mulatta): a pilot study. J Comp Psychol. 1999;113:59.

43. Shutt K, MacLarnon A, Heistermann M, Semple S. Grooming in Barbary macaques: better to give than to receive? Biol Letters. 2007;3:231–233.

44. Akinyi MY, Tung J, Jeneby M, Patel NB, Altmann J, Alberts SC. Role of grooming in reducing tick load in wild baboons (Papio cynocephalus). Anim Behav. 2013;85:559–568.

45. Rosengaus RB, Traniello JFA, Lefebvre ML, Carlock DM. The social transmission of disease between adult male and female reproductives of the dampwood termite Zootermopsis angusticollis. Ethol Ecol Evol. 2000;12:419–433.

46. Nunn C, Altizer S, Altizer SM. Infectious diseases in primates: behavior, ecology and evolution. Oxford University Press; 2006 Apr 27.

47. Huffman MA, Chapman CA. Primate parasite ecology: the dynamics and study of host-parasite relationships. 2009.

48. Dowling DK, Richardson DS, Komdeur J. No effects of a feather mite on body condition, survivorship, or grooming behaviour in the Seychelles warbler, Acrocephalus sechellensis. Behav Ecol Sociobiol. 2001;50:257–262.

49. Brain C, Bohrmann R. Tick infestation of baboons (Papio ursinus) in the Namib Desert. J Wildlife Dis. 1992;28:188–191.

50. Townsend AK, Hawley DM, Stephenson JF, Williams KE. Emerging infectious disease and the challenges of social distancing in human and non-human animals. Proceedings of the Royal Society B. 2020 Aug 12;287(1932):20201039.

51. Drewe JA. Who infects whom? Social networks and tuberculosis transmission in wild meerkats. P Roy Soc Lond B Bio. 2010;277:633–642.

52. Konrad M, Vyleta ML, Theis FJ, Stock M, Tragust S, Klatt M, Drescher V, Marr C, Ugelvig LV, Cremer S. Social transfer of pathogenic fungus promotes active immunisation in ant colonies. PLOS Biol. 2012;10:e1001300.

53. Filmalter N. A vegetation classification and management plan for the Hondekraal section of the Loskopdam Nature Reserve [dissertation]. Pretoria: University of South Africa; 2010.

54. Barrett AS, Brown LR, Barrett L, Henzi P. A floristic description and utilisation of two home ranges by vervet monkeys in Loskop Dam Nature Reserve, South Africa. Koedoe. 2010;52:1–12.

55. Wren BT. Behavioural ecology of primate-parasite interactions [dissertation]. West Lafayette (IN): Purdue University; 2013.

56. Isbell LA, Young TP. Social and ecological influences on activity budgets of vervet monkeys, and their implications for group living. Behav Ecol Sociobiol. 1993;32:377–385.

57. Fruteau C, Lemoine S, Hellard E, Van Damme E, Noë R. When females trade grooming for grooming: testing partner control and partner choice models of cooperation in two primate species. Anim Behav. 2011;81:1223–1230.

58. Fruteau C, Voelkl B, Van Damme E, Noë R. Supply and demand determine the market value of food providers in wild vervet monkeys. P Natl Acad Sci. 2009;106:12007–12012.

59. van de Waal E, Renevey N, Favre CM, Bshary R. Selective attention to philopatric models causes directed social learning in wild vervet monkeys. P Roy Soc Lond B Biol. 2010;rspb20092260.

60. Pansini R. Induced cooperation to access a shareable reward increases the hierarchical segregation of wild vervet monkeys. PLOS One. 2011;6:e21993.

61. van de Waal E, Krützen M, Hula J, Goudet J, Bshary R. Similarity in food cleaning techniques within matrilines in wild vervet monkeys. PLOS One. 2012;7:e35694.

62. Wren BT, Gillespie TR, Camp JW, Remis MJ. Helminths of Vervet Monkeys, Chlorocebus aethiops, from Loskop Dam Nature Reserve, South Africa. Comp Parasitol. 2015;82:101–108.

63. Martin P, Bateson P. Measuring behaviour: an introductory guide. 3rd ed. Cambridge: Cambridge University Press; 2007.

64. Gillespie TR. Noninvasive assessment of gastrointestinal parasite infections in free-ranging primates. Int J Primatol. 2006;27:1129–1143.

65. Gillespie TR, Morgan D, Deutsch JC, Kuhlenschmidt MS, Salzer JS, Cameron K, Reed T, Sanz C. A legacy of low-impact logging does not elevate prevalence of potentially pathogenic protozoa in free-ranging gorillas and chimpanzees in the Republic of Congo: logging and parasitism in African apes. EcoHealth. 2009;6:557–564.

66. Bush AO, Lafferty KD, Lotz JM, Shostak AW. Parasitology meets ecology on its own terms: Margolis et al. revisited. J Parasitol. 1997;575–583.

67. Ghai RR, Fugere V, Chapman CA, Goldberg TL, Davies TJ. Sickness behaviour associated with non-lethal infections in wild primates. P Roy Soc B Biol. 2015;282:20151436.

68. Stephenson LS, Holland CV, Cooper ES. The public health significance of Trichuris trichiura. Parasitology. 2000;121:S73–S95.

69. Holmes JC, Bethel WM. Modification of intermediate host behaviour by parasites. Behav Aspects Parasite Transmission. 1972;51:123–149.

70. Poulin R. Host specificity. In: Evolutionary ecology of parasites, 2nd ed. London: Chapman Hall; 1998. p. 41–69.

71. Moore J. Parasites and the behaviour of animals. Oxford: Oxford University Press; 2002.

72. Poulin R. Parasite manipulation of host behaviour: an update and frequently asked questions. Adv Stud Behav. 2010;41:151–186.

73. Webster JP. The effect of Toxoplasma gondii on animal behaviour: playing cat and mouse. Schizophrenia Bull. 2007;33:752–756.

74. Joshua K, Yidawi JP, Sada A, Msheliza EG, Turaki UA. Prevalence and morphotype diversity of Trichuris species and other soil-transmitted helminths in captive non-human primates in northern Nigeria. J Threat Tax. 2020;12(10):16239–44.

75. Egbetade A, Akinkuotu O, Jayeola O, Niniola A, Emmanuel N, Olugbogi E, Onadeko S. Gastrointestinal helminths of resident wildlife at the Federal University of Agriculture Zoological Park, Abeokuta. Sokoto J Vet Sci. 2014;12(3):26–31.

76. Hart BL. Biological basis of the behaviour of sick animals. Neurosci Biobehav R. 1988;12:123–137.

77. Krief S, Huffman MA, Sévenet T, Guillot J, Bories C, Hladik CM, Wrangham RW. Noninvasive monitoring of the health of Pan troglodytes schweinfurthii in the Kibale National Park, Uganda. Int J Primatol. 2005;26:467–490.

78. Barber I, Hoare D, Krause J. Effects of parasites on fish behaviour: a review and evolutionary perspective. Rev Fish Biol Fisher. 2000;10:131–65.

